# A Novel Art of Electrocardiogram Assessment in Zebrafish for Cardiovascular Disease Studies and Drug Screening

**DOI:** 10.1101/2021.07.02.450921

**Authors:** Tai Le, Jimmy Zhang, Anh Hung Nguyen, Ramses Seferino Trigo Torres, Khuong Vo, Nikil Dutt, Juhyun Lee, Yonghe Ding, Xiaolei Xu, Michael P.H. Lau, Hung Cao

**Affiliations:** Department of Electrical Engineering and Computer Science, UC Irvine, Irvine, CA, 92697, USA; Department of Biomedical Engineering, UC Irvine, Irvine, CA, 92697, USA; Donald Bren School of Information and Computer Sciences, UC Irvine, CA 92697, USA; Department of Bioengineering, University of Texas, Arlington, TX, 76019, USA; Department of Biochemistry and Molecular Biology, Mayo Clinic, Rochester, Minnesota, 55905, USA; Sensoriis., Inc, Edmonds, WA, 98026, USA

## Abstract

The zebrafish (Dario rerio) has proven to be an excellent animal model for biological research owing to its small size, low cost for maintenance, short generation time, amenable genetics, and optical transparency. Zebrafish have been extensively used in cardiovascular studies in which mutant lines with cardiovascular defects were introduced and analyzed. Despite the small size, technological advances have paved the way to effectively assess cardiac functions of zebrafish. Here, we present a novel art for long-term simultaneous monitoring and analysis of electrocardiogram (ECG) in multiple zebrafish with controlled environment. The system helps minimize the effect of anesthetic drug and temperature to cardiac rhythm side effects as well as save time and efforts by 40-50 fold compared with the conventional approach. We further employed the system to study the Na^+^ sensitivity in the development of sinus arrest in *Tg(SCN5A-D1275N)* fish, a study model of the sick sinus syndrome, as well as the relationship between this variant and drug administration. The novel ECG system developed in this study holds promise to greatly accelerate other cardiovascular studies and drug screening using zebrafish.

## Introduction

Cardiovascular diseases (CVDs) are the leading cause of death worldwide. According to the 2020 AHA Annual Report, almost 860,000 people died of CVDs in the U.S. in 2017, and the overall financial burden from CVDs totaled $351.2 billion in 2014-2015, emphasizing the urgency to explicate the etiologies of CVDs [1]. One such CVD is the sick sinus syndrome (SSS), a collection of progressive disorders marked by the heart’s inability to maintain a consistent rhythm of heart muscle contraction and relaxation [2]. It is characterized by age-associated dysfunction of the sinoatrial node (SAN), with varying symptoms such as syncope, heart palpitations, and insomnia [3]. The SSS has multiple manifestations on electrocardiogram (ECG) recordings, including sinus bradycardia, sinus arrest, and sinoatrial block. The pathophysiology of SSS is not fully understood, but research has determined that it can be caused by numerous factors ranging from pharmacological medications and sleep disturbances to fibrosis and ion channel dysfunction [4]. The current primary treatment for SSS is the implantation of an artificial pacemaker. While pacemakers [5] have been effective in treating various arrhythmic conditions [6], they also carry their own complications. Since the pacemaker is seen as a permanent solution, long-term issues such as lead displacement, hematomas, and pneumothorax are significant [7], and conditions such as thromboembolism and stroke are documented [8, 9]. Additionally, because of the multifactorial nature of SSS, new arrhythmic symptoms could arise after pacemaker implantation, as seen in pacemaker syndrome [10]. All these issues could worsen a patient’s condition after pacemaker implantation. As a result, permanent alternative solutions should be investigated, including utilizing a biological means to naturally regenerate or replicate the function of the SAN. Previous research has emphasized SSS-associated genetic pathways as potential avenues to a more permanent treatment for SSS. Gene therapy techniques have been utilized to rescue SAN function. Overexpression of *Tbx18* and *Tbx3* via cardiomyocyte transfection has been shown to develop cells with SAN-like function, forming a biologically produced pacemaker. These genes were linked to the expression of connexin proteins, such as *Cx40*, and *Cx43*, which were crucial in maintaining the proper propagation of cardiac conduction [11–13]. Several animal models (*e.g.* mice, rats, pigs) have been tested with this strategy, which have resulted in generally positive results [14–17]. However, gene therapy does have some limitations such as its temporary nature and heterogeneity (not all cells are transfected). Additionally, no study has demonstrated its efficacy in humans, as all studies were conducted in animal models. Therefore, more research should be conducted to identify the electrophysiological phenotypes of these genetic anomalies to help devise future treatment methods.

Zebrafish serve as an ideal model for cardiovascular studies because of their similar homology to humans in both morphology, physiology, and genetics. Despite having only two discernible chambers in the zebrafish heart compared to four in human hearts, the zebrafish heart possesses a similar contractile structure with an analogous conduction system [18, 19]. The zebrafish genome also carries remarkable homology to that in humans, as 70% of all human genes have orthologues in the zebrafish genome, making the zebrafish model an applicable model in studying genetic pathways [20]. Therefore, the zebrafish model is appropriate in the study of SSS and the correlation of related genetic pathways to the electrophysical phenotype via ECG. Currently, several research groups have developed zebrafish ECG acquisition and sensors to support cardiac studies. Regarding sensor design, needle electrodes are the most commonly used. Lin *et al.*, [21] designed and tested the needle electrode with different materials, including tungsten filament, stainless steel and silver wire to investigate the recorded signal quality. Along with a portable ECG kit, the authors aim to provide a standard platform for research and teaching laboratories. The needle system is also deployed in other studies [21–24] to conduct biological and/or drug-induced research. All needle-based studies demonstrated promising results; however, to collect favorable signals, the needles need to be gently inserted through the dermis of zebrafish. This could cause injury to the fish’s heart, thus possibly changing signal morphology [23]. Moreover, it requires an intensive effort in precisely positioning the electrode on the heart to achieve a clear ECG signal. Therefore, several alternative probe systems have been developed, including the micro-electrode array (MEA) and the 3D-printed sensors. Our team and others have demonstrated the use of MEA for acquisition and provided the signal with favorable signal-to-noise ratio (SNR) with high spatial and temporal resolution [25–27]. For instance, we presented a MEA array covering the fish’s heart which enables site-specific signals [26, 27]. Cho *et al.*, [28] developed a MEA printed on a flexible printed circuit board (FPCB) based on a polyimide film for multiple electroencephalogram (EEG) recording for epilepsy studies. Although the MEA allows multiple signal recordings, only one fish can be assessed at a time due to the limiting number of channels on acquisition. Moreover, most studies have used bulky and expensive acquisition tools to collect data and then transfer them through a cable to a computer. These require a designated benchtop area to conduct experiments, yet potentially encounters errors due to instability of cable and connectivity. In the market, commercially available systems, *e.g.* the one from iWORX (Dover, NH) with a compact amplifier, can improve the mobility of the system. However, several challenges have not been resolved: **i**) the current systems only record for a short period of time, which result in inconsistent results among different fish; **ii**) the ECG acquisition requires anesthetized animals, rendering it stressful to the fish and inadequate to provide intrinsic cardiac electrophysiological signals; **iii**) manual one-by-one measurement limits the ability of doing studies required to test with a large number of fish; and **iv**) signal processing has been carried out offline with humongous efforts.

In this work, we introduce a novel system, Zebra II, capable of obtaining long-term ECG recordings for multiple fish simultaneously. An in-house electronic device is developed, leveraging the Internet of Thing (IoT) capability with wireless data transmission and data processing on a mobile application. This enhances the mobility and versatility of the system to conduct research on zebrafish models. The system is validated through numerous experiments, showing its potential with 1) simultaneous measurement for up to 4 fish; 2) continuous ECG recording for up to 1 hour compared to several minutes of other systems; 3) reduction in arrhythmic side effects with the use of 50% lower Tricaine concentration. Furthermore, we investigate a specific electrophysiological phenotype, namely sinus arrest, induced by sodium chloride on mutant fish *Tg(SCN5A-D1275N)* to demonstrate that our proposed system can be used as a screening tool to detect and elucidate zebrafish cardiac arrhythmic symptoms.

## Results

### Demonstration of Zebra II for multiple zebrafish ECG recording

The Zebra II system is designed to allow the fish to stay comfortably for up to 1 hour while the ECG signal is acquired. It comprises of a perfusion system, soft polymer housings, sensors and an electronic system (**Fig. 1a**). The perfusion system continuously provides the low concentration of anesthetic drug (MS-222), reducing the aggressiveness and activity of zebrafish and giving the fish with adequate oxygen levels. Thus, with apparatuses made of polydimethylsiloxane (PDMS), the perfusion system can keep them stable for a long time during the measurement. Moreover, a home-made thermo box is designed with a thermostat control, a light bulb, and a temperature sensor, which automatically controls the temperature of environment to conduct experiments. An in-house electronic system and a mobile application are fully developed, allowing wireless data transmission as well as data storage and analysis, thereby greatly reducing time and effort. (**Fig. 1b-d, Sup. Fig. 1**). The overall fish ECG system specification is shown in **Sup. Table 1**. Finally, the data are wirelessly transmitted to a mobile application as shown in **Fig. 1d**. The mobile app allows real-time data transmission and send the data to the cloud system. The data transmission is further illustrated in **Sup. Fig. 2**.

**Figure 1.**
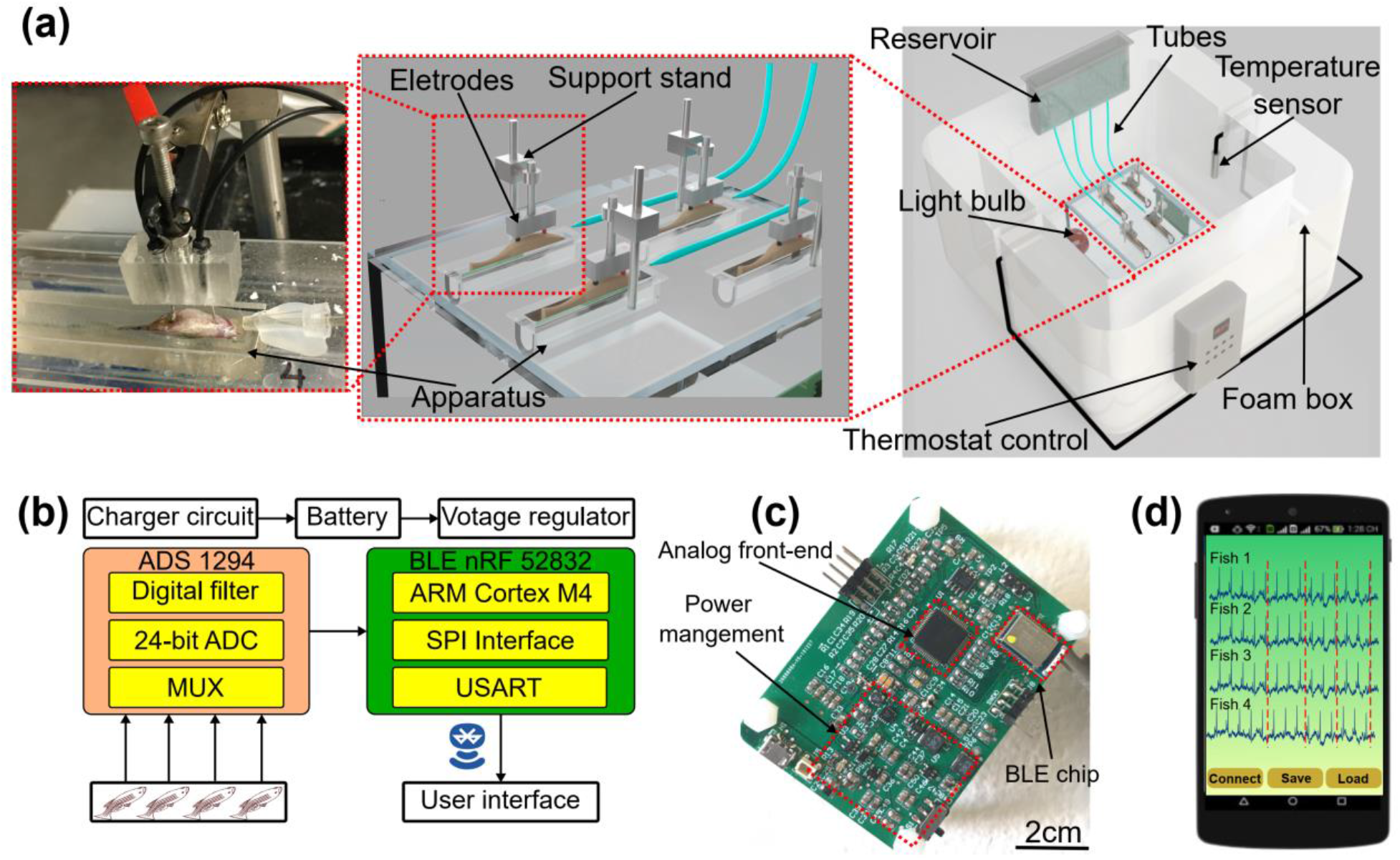
Design of the prolonged ECG system for multiple adult zebrafish recording. (**a**) the prolonged ECG mechanical design: the reservoir containing solution, the tube system dropping the solution on the fish, the electrode and support stand recording the ECG signal. (**b**) System-level block diagram showing analog front-end chip, signal transduction, wireless transmission for the ECG signal to user interface. (**c**) In-house electronic board having system-on-chip for wireless transmission, power management connecting to the electrode for ECG acquisition. (**d**) User interface of mobile application receiving ECG signal from multiple fish.

Numerous experiments were conducted to validate the performance of Zebra II. First, the zebrafish ECG was collected simultaneously by the Zebra II and a commercial device developed by iWORX (Dover, NH). The signals were then compared in both frequency domain and time domain (**Sup. Fig. 3a&b**). Specifically, the correlation coefficients were 98.78% and 96.54% in time domain and frequency domain, respectively. Moreover, the heart rate value and QRS interval were also compared (**Sup. Fig. 3c&d**). As seen, the Zebra II’s performance is comparable to that of the commercial iWORX device. Further, we performed another experiment on 36 wildtype (WT) zebrafish dividing to 2 groups: 1) control (n = 20) and 2) Amiodarone treated (n = 16) fish. Amiodarone is used to prevent various types of arrhythmic symptoms, including ventricular tachycardia and atrial fibrillation [29]. However, Amiodarone has been also reported to cause bradycardia and to prolong the QT interval in zebrafish [30]. This experiment allowed us to assess the drug screening capacity of the Zebra II system. Specifically, for the treated group, fish were treated with 100 μM Amiodarone by immersing them in a tank with 200 mL of the Amiodarone medium. ECG was recorded after 1 hour of immersion. This experiment was conducted with both our Zebra II and iWORX systems. As shown in **Sup. Fig. 4**, heart rate (HR) value, QRS duration and QTc interval were analyzed. With the control group, there is no significant difference (p-value > 0.05) between two systems in terms of QRS and QTc value. Similarly, the HR value and QTc value show no significant difference in treated group. Furthermore, the Bland Altman analysis in **Sup. Fig. 4b, d, f** shows the agreement level between two systems with most of HR values and QTc values belonged to the limit of agreement (LOA) region.

### Investigation of tricaine and temperature to reduce cardiac rhythm side effect

As shown in **Fig. 2a**, the ECG morphology was observed with different doses. The ECG signal shows gill motion noise, interfering ECG waveforms such as P, T and QRS waves with 75 ppm Tricaine while the signal appears to be more stable under 100 ppm and 150 ppm treatment, providing clear ECG waves. After 40-min long measurement, the recovered time and survival rate of fish are collected (**Fig. 2b**). It was found that fish under higher Tricaine concentration need longer recovery time. Specifically, it takes the average of 7 minutes to recover the fish under 150 ppm while those under 75 ppm and 100 ppm treatment it requires 3 minutes and around 4.2 minutes for fish to wake up after measurement, respectively. Furthermore, with 150 ppm treatment, the survival rate is about 75% while other concentrations can keep the fish alive after measurement above 90%. It reflects to the effect of a high dose used under a long period of time measurement which is similar to the dose for fish euthanasia (*i.e.*, 168 ppm) [31]. Given the recovery time and survival rate along with ECG morphology collected for different Tricaine concentrations, the dose of 100 ppm is the optimal one for the prolonged measurement. Regarding the heart rate variation every 5 minutes during the measurement, we characterized it with the ECG signal under 75 ppm and 100 ppm (**Sup. Fig. 5**). After 100 ppm treatment, the heart rate variation shows no significant difference among first 30 minutes, making the average standard deviation (STD) of 17 beats per minute (BPM). In contrast, that value with the ECG data under 75 ppm describes the changes among every 5-minute data, which leads to 22 BPM difference based on STD.

**Figure. 2.**
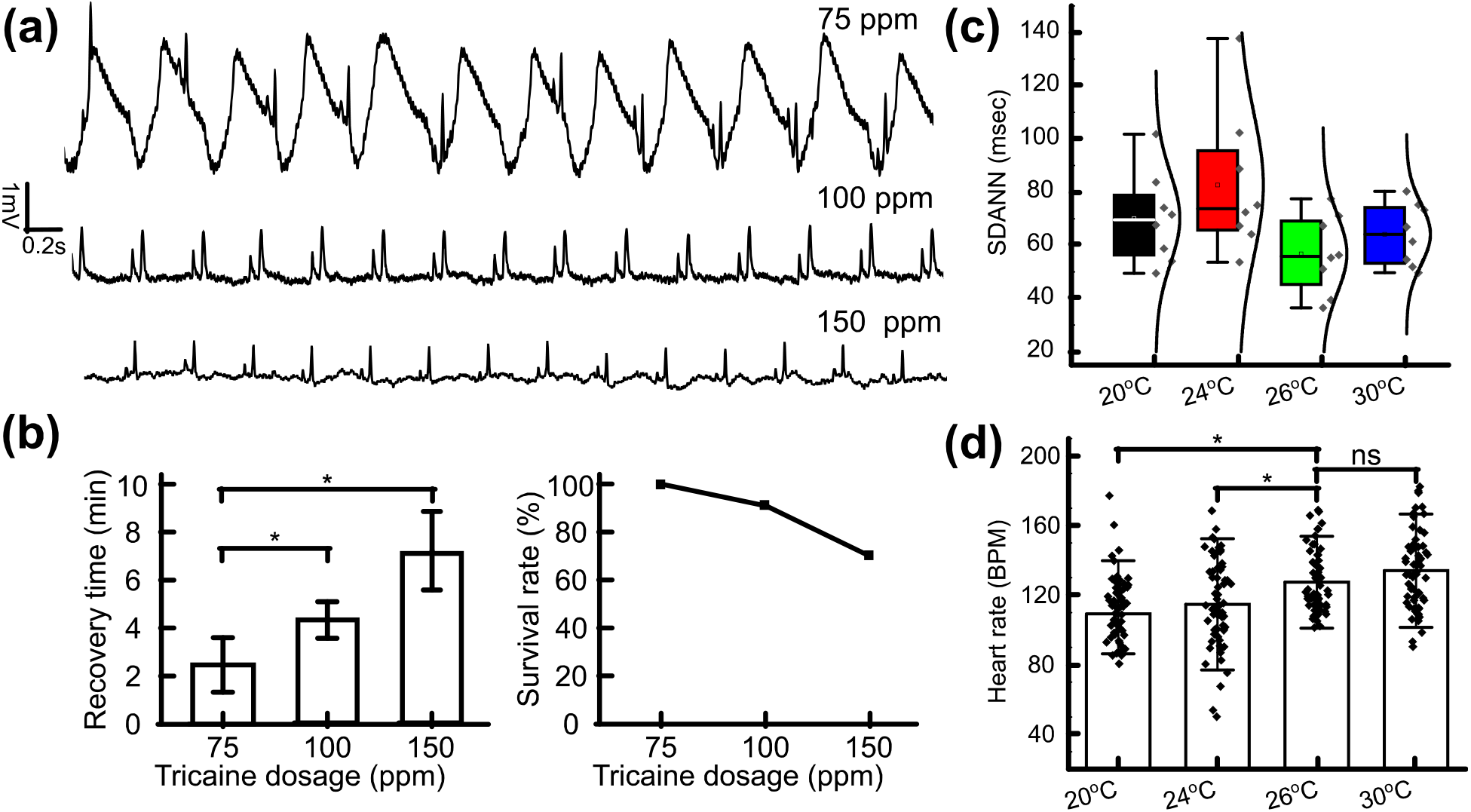
Investigation of tricaine and temperature to reduce cardiac rhythm side effect. (**a**) ECG morphology example recorded by different Tricaine concentrations. (**b**) Bar chart comparing recovery time needed after treatment for each Tricaine concentration. Line graph describing the survival rate of zebrafish treated by different Tricaine concentrations. (**c**) SDANN in WT fish with different temperatures. (d) Heart rate in WT fish with different temperatures. *p <0.05; **p < 0.01 (one-way analysis of variance). ns indicates not significant.

Given the optimal Tricaine concentration, different temperatures are investigated. A temperature-control incubator was designed for the experiment. The zebrafish ECG system was put in the middle of the incubator. Temperature within the chamber could range from 20°C to 32°C, as measured by a thermometer inside the chamber and controlled by a thermostat with accuracy of ±1°C. Prior to recording the ECG signals in this experiment, the impedance of the electrodes was measured on zebrafish skin (**Sup. Fig. 6**) ensuring the signal stability during the long-term measurement. As shown in **Fig. 2c**, the SDANN at 26°C has the lowest value with the range of 36 msec to 75 msec while that at 24°C is highest with the range of 50 msec to 139 msec. Moreover, the data distribution from heart rate value collected by every 5 minutes collected at 26°C is the most condensed (**Fig. 2d**). Thus, under 26°C, the heart rate is more stable than that under other temperatures.

### Response analysis to drug treatment in real time with the Zebra II system

As shown in **Fig. 3a**, 4 fish were measured simultaneously, and 4 doses were consecutively filled in the reservoir and each dose lasted around 5 minutes. The change in response to different dosages in all four fish was obvious. Zooming out the data collected from fish 1 in different amiodarone dosages as denoted from (1) to (4), the QTc interval showed considerable changes. For instance, the QTc value is 310 msec without drug treatment and it tends to increase after the fish got treated. Specifically, it is 330 msec with 70 μM of amiodarone while it is 476 msec and 536 msec with 100 μM and 200 μM of amiodarone, respectively. **Fig. 3b** describes the overall changes in terms of QT prolongation, QRS interval and HR value in response to different amiodarone concentration. With the increase of amiodarone dosage, QT prolongation and QRS interval shows an increase while the average heart rate is decreased.

**Figure 3.**
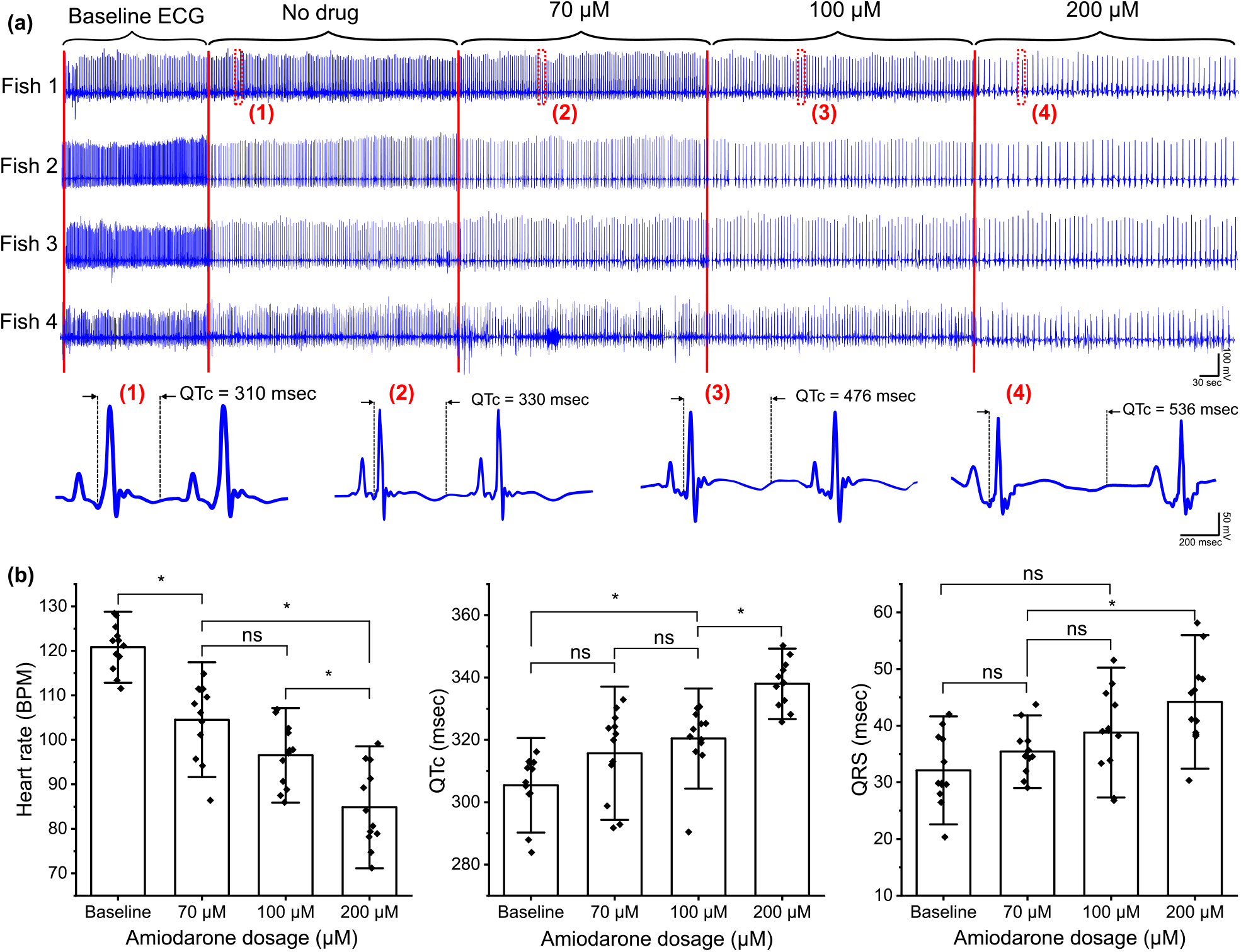
Demonstration of the prolonged ECG system showing the ECG morphology in response to different amiodarone concentrations. (**a**) The representative of ECG signal recorded by the proposed system and its change of signal morphology due to different amiodarone dosages in real time. (**b**) Bar chart describing the discrepancy of HR value, QTc interval and QRS interval in ECG signal with different amiodarone dosages (n = 8 fish). *p <0.05 (one-way analysis of variance with Turkey test). ns indicates not significant.

### Evaluation of Na+ sensitivity in the development of sinus arrest (SA) in *Tg(SCN5A-D1275N)*

WT fish (n = 12, aged 1.5 years) and *Tg(SCN5A-D1275N)* fish (n = 8, aged 10 months) were used in this experiment. **Fig. 4a** illustrates the ECG morphology of *Tg(SCN5A-D1275N)* with different NaCl dosages. We noticed that with a small NaCl dosage (0.1 ‰), the zebrafish starts showing the reduction in heart rate, followed by significant decrease in higher dosages. According to the SA criteria (*i.e.*, RR interval is greater than 1.5 sec) determined in our previous work [32], sinus arrest appears more frequently after treatment with 0.6 ‰ NaCl and above (**Table 1**). As shown in **Fig. 4b**, the HR value started significantly dropping in 0.6 ‰ NaCl treatment in the SSS mutant. In contrast, NaCl does not show profound effect to the control group (WT fish) as evidenced by the slight decrease in the HR responding to different NaCl levels. It was worth noting that these WT fish were at 1.5 years old, which could attribute to an increase of SA [32], causing the slight reduction of HR in the experiment (**Sup. Fig. 7**). In terms of heart rate variability (HRV), *Tg(SCN5A-D1275N)* fish showed a remarkable increase with the high NaCl dosages (0.9 and 1.8 ‰) compared with other dosages. This provides evidence that the *Tg*(*SNC5A-D1275N*) triggers more sinus arrest under NaCl treatment (**Table 1**). Moreover, QTc interval was also characterized and it shows the same pattern happened in SDNN with the increase of QTc in response to NaCl dosages (**Fig. 4d, Sup. Fig. 8**).

**Figure 4.**
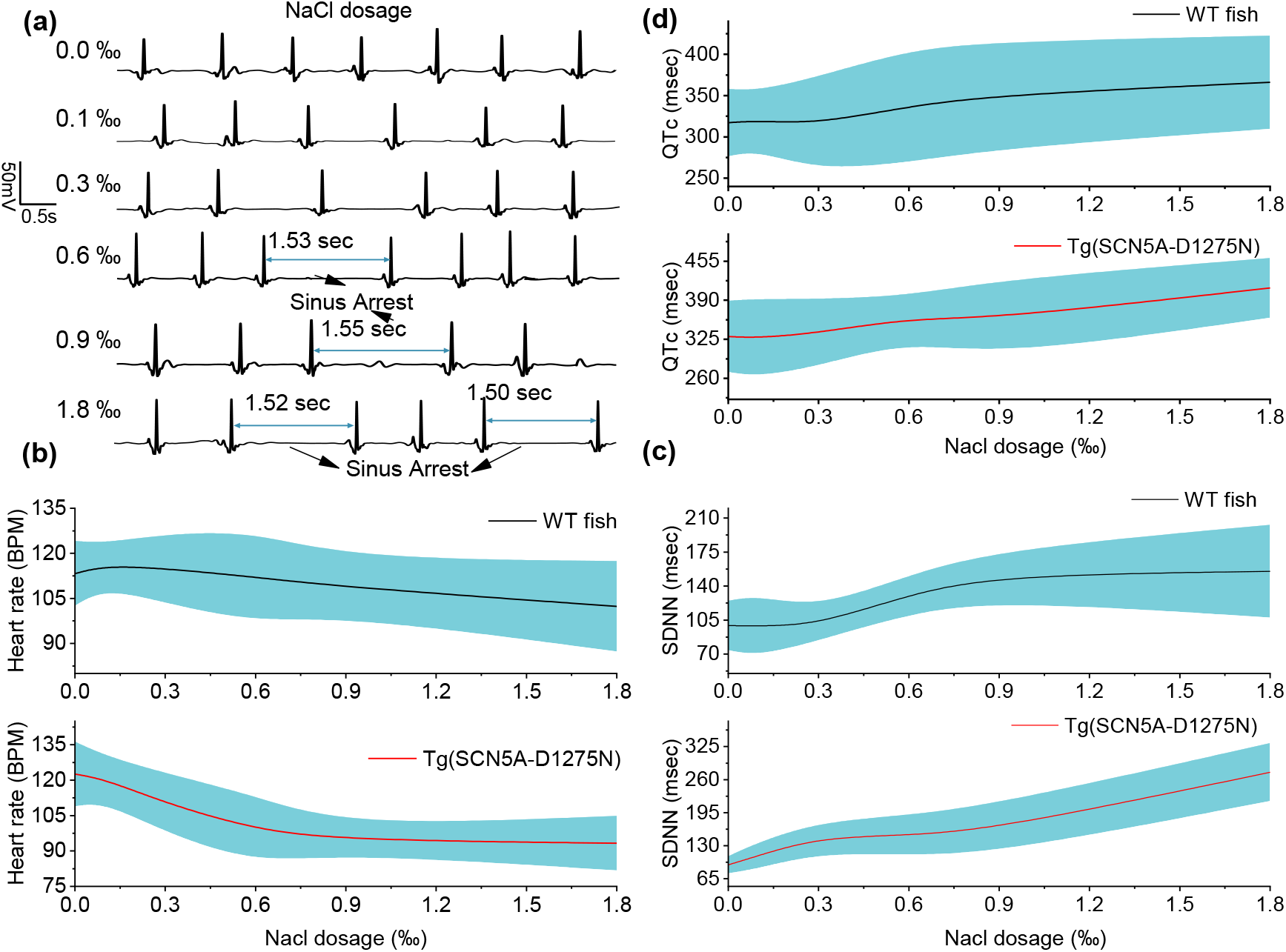
Evaluation of Na+ sensitivity in the development of sinus arrest in *Tg*(*SCN5A-D1275N*). (**a**) the representative ECG waveforms before and after NaCl treatment with different dosages. The sinus arrest is to be appear more in response to the increase of the NaCl dosage. (**b**) the average heart rate of wild-type fish (n = 12) and mutant fish (n = 8) with each dosage of NaCl. (**c**) Standard deviation of normal-normal beat sinus (SDNN) of wild-type fish and mutant fish. (**d**) QTc values of two types of fish after treatment with different NaCl concentration.

**Table I.**

| Fish | WT |  |  |  | Tg(*SCN5A-D1275N*) |  |  |  |
| --- | --- | --- | --- | --- | --- | --- | --- | --- |
| Drug | Average HR (BPM) | Percentage of Fish with SA (No. of cycles) | SA frequency epm | QTc (msec) | Average HR (BPM) | Percentage of Fish with SA (No. of cycles) | SA frequency epm | QTc (msec) |
| 0 ‰ NaCl | 113.3 ± 10.8 | 8.3(1) | 0.08 | 317.2 ± 40.7 | 122.6 ± 13.7 | 12.5 (1) | 0.125 | 329.8 ±59.1 |
| 0.1‰ NaCl | 115.9 ± 7.5 | 8.3(1) | 0.08 | 319.7 ± 36.3 | 120.4 ± 9.9 | 12.5(1) | 0.125 | 327.2 ±64.8 |
| 0.3‰ NaCl | 114.9 ± 11.5 | 8.3(1) | 0.08 | 316.1 ± 56.6 | 110.3 ± 12.6 | 37.5(7) | 0.875 | 335.7 ±55.8 |
| 0.6‰ NaCl | 112.1 ± 15.2 | 16.6(2) | 0.16 | 337.2 ± 68.2 | 98.9 ± 14.4 | 75(8) | 1 | 359.9 ±37.1 |
| 0.9‰ NaCl | 108.7 ± 9.5 | 25(3) | 0.25 | 350.7 ± 64.7 | 94.6 ± 5.7 | 75(15) | 1.875 | 360.1 ± 64.0 |
| 1.8‰ NaCl | 102.4 ± 15.0 | 50(7) | 0.42 | 366.1 ± 56.4 | 93.3 ± 11.5 | 87.5(17) | 2.125 | 410.5 ±49.5 |
| 0.9‰ NaCl + 50 µM Meth | 109.4 ± 12.3 | 25(3) | 0.25 | 360.2 ± 70.3 | 96.95 ± 21.2 | 75(15) | 1.875 | 391.3 ± 76.1 |

As mentioned previously, the sodium channel gene *SCN5A* is one of the most frequently mutated genes in LQTS. To further investigate this variant as a candidate for cardiac studies, we assess the relationship between *SCN5A* and methamphetamine (Meth) – a controlled substance. Several groups have studied its connection of using addictive drugs with sudden death. For instance, Sayaka, *et al.*, screened several variations in the LQTS-associated genes *KCNQ1 (LQT1)* and *KCNH2 (LQT2)* showing the link to the risk of serious cardiac arrhythmia for those abusing addictive drugs [33]. However, the authors do not test for *SCN5A* variants which provided us an opportunity to explore its effect. Additionally, we sought to rescue the bradycardic symptoms induced by NaCl to provide insight on the nature of those symptoms, as Meth has been previously demonstrated to increase heart rate after administration [34]. Specifically, we treated two groups (control – WT fish and treated – *Tg(SCN5A-D1275N)*) in 0.9 ‰ NaCl in 30 minutes before immerging the fish in 50 μM Meth in other 30 minutes. As shown in **Fig. 5**, the HR value, SDNN and QTc value were compared between two groups with three critical moments, including without drug treatment, with NaCl treatment and combined NaCl and Meth treatment (n = 12 WT fish and n = 8 *Tg*(*SCN5A-D1275N*)). The average heart rate after the combined NaCl and Meth treatment (96.96 ± 7.61 BPM) showed a slight increase compared with that solely treated with NaCl (94.59 ±5.69 BPM) with *Tg(SCN5A-D1275N)*; however, it showed no significant difference between two moments (P > 0.05). Similarly, the SDNN value does not show any significant difference. This means the Meth treatment did not affect the HR and SDNN in both groups (**Table 1**), implying that NaCl administration resulted in irreversible bradycardic symptoms that could not be easily treated with agents that increase heart rate. In contrast, we found a significant increase of QTc interval in the *Tg(SCN5A-D1275N)* group before (360.1 ± 64.0 msec) and after (391.3 ± 76.1 msec) Meth treatment, indicating the additive effect of Meth to QT prolongation (**Table 1**- in red).

**Figure 5.**
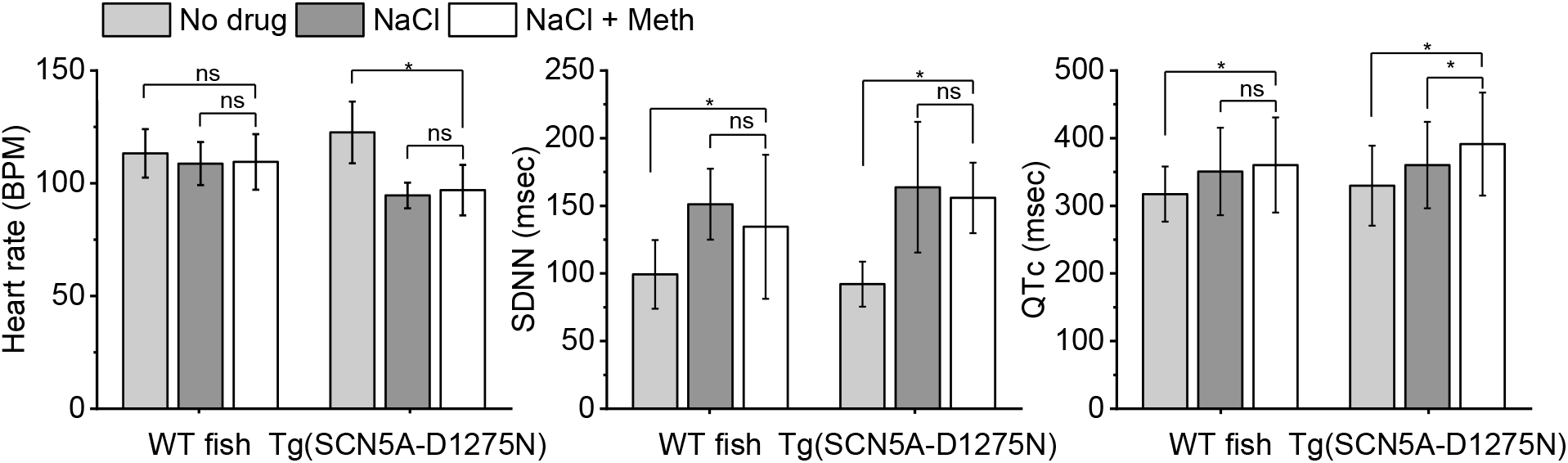
Investigation of methamphetamine (Meth)’s efficacy to rescue the heartbeat sinus after treatment with NaCl. The experiment was analyzed and compared heart rate, QTc and SDNN among three stages (i.e., no drug treatment, with NaCl and with NaCl + Meth) and the experiment was conducted in both WT fish and mutant fish Tg(*SCN5A-D1275N*).

## Discussion and Conclusion

A strong association between high sodium intake and cardiovascular disease has been reported in hypertensive populations [35]. A high sodium diet is associated with alterations in various proteins responsible for transmembrane ions homeostasis and myocardial contractility. Recent studies provided important evidence that excess sodium promotes structural and functional impairment of the heart, especially in populations bearing mutant phenotypes of the major cardiac sodium channels. These mutations are responsible for various types of cardiac disorders, including Brugada syndrome (*BrS*), long QT syndrome (*LQT3*), cardiac conduction disease (CCD), sick sinus syndrome (SSS), atrial fibrillation (AF), progressive cardiac conduction defect (PCCD) [36]. In the original reports [36–39], *SCN5A* mutations are associated with cardiac conduction defect and atrial arrhythmias which cause bradycardia with reduced heart rate. In the present study, the developed system shows the susceptible effects of excess sodium ions to abnormal cardiac rhythm of the *Tg(SCN5A-D1275N)* mutant, to validate our system with different arrhythmic phenotypes. Our noticeable finding is that the excessive sodium ions cause sinus arrest in *Tg(SCN5A-D1275N)* at 0.6 ‰, 0.9 ‰, and 1.8 ‰, corresponding to 1.53 sec, 1.55 sec and 1.52 sec, respectively; and slower heart rate and prolonged QTc are observed only in mutant fish. These results provide a significant association between the increased frequency of sinus arrest, slower heart rate, and prolonged QTc with increased sodium intake in SSS mutants. According to previous reports [40, 41], the *SCN5A* sodium-channel protein can disrupt the heart’s electrical activity and lead to a dramatic decrease of heart rate. The slow-conducting *Tg(SCN5A-D1275N)* phenotype has been proved by voltage-clamp measurement [42, 43] in which data were consistent with our finding. In the eight fish *D1275N* carriers, the average QTc intervals were 385 msec at the upper physiological limit, indicating that the QTc intervals in *Tg(SCN5A-D1275N)* fish are generally more prolonged than wild type animals. Moreover, excess Na^+^ ions cause not only slow heart rate and prolonged QTc but also increased sinus arrest frequency in *Tg(SCN5A-D1275N)* (**Fig. 4**).

Sodium-overload sinus arrest observed in this study may be associated with a rise of the intracellular Na+ in heart muscle due to gain-of-function of *Tg(SCN5A-D1275N)* for sodium ions traveling into the cell. Detection of Na^+^-induced sinus arrest by the developed system shows that Brugada syndrome mutation *Tg(SCN5A-D1275N)* is susceptible to excess Na^+^ ions due to hastening epicardial repolarization and causing idiopathic ventricular conduction, which induce ECG changes and ventricular arrhythmias of Brugada syndrome. Moreover, the mutation (*D1275N*) evokes the long QT syndrome which is caused by excessive INa detected by the developed system with continuously prolonged measurement. Recorded data by the developed system is consistent with clinical reports indicating that Brugada syndrome in human and animals have reported Na^+^-induced abnormalities in ventricular conduction [44, 45]. Thus, an overload of Na^+^ ions can cause destabilized closed-state inactivation gating of *D1275N* that may attenuate the ventricular conduction delay, shown in arrhythmic parameters (**Table 1**).

One of the key novelties of the Zebra II system is the capacity to test multiple drugs on one fish with a continuously prolonged assay. The analysis of ECG of *Tg(SCN5A-D1275N)* indicates that Methamphetamine do not improve sinus arrest frequency and heart rate. However, it caused prolonged QT at 50 μM of Meth. In both groups, the QTc was longer (by 350 msec for wild type and 385 msec for *Tg(SCN5A-D1275N)* after Methamphetamine treatment. Thus, the robust performance of the system allows incorporation the multiple drugs with different effects (*e.g.*, antagonistic effects) in a single continuously prolonged assay to study drug-drug interactions on a specific arrhythmic phenotype, which is heavily performed by the short time course of current systems. Extending ECG measurement in merely-sedated fish allows measuring interactive effects of different drugs on a specific phenotype by a prolonged screening course. Different ECG phenotypes are recorded using the prolonged real-time courses (over 40 min) to provide intuitive insights into how the drug interaction effects, indicating a tool to evaluate drug efficacy. As shown in **Fig. 4d**, the average SDNN, the standard deviation of normal to normal R-R intervals, was 125 msec for wild type fish, but was 255 msec for *Tg*(*SCN5A-D1275N*), consistent with reduced conduction velocities due to Na^+^ ion channel disfunction [46].

The number of conduction defects associated with the *D1275N* mutation provide a biophysiological mechanism for conduction defects observed in *Tg(SCN5A-D1275N)* and analyzed by the developed system. The *D1275N* mutant channels provides new evidence that excess Na^+^ ions in mutant sodium channel dysfunction can produced isolated conduction disease, with pathological slowing of the heart rate, prolonged QTc and higher frequency of sinus arrest. Although functional data are not available for the system performance, consistent data of conduction disease and Na^+^ channel function in real-time and prolonged continuously measurement indicates that the developed system can be used for many applications including drug screening and interactions with various zebrafish mutant phenotypes.

In order to enable remote monitoring and high-throughput zebrafish ECG analysis, our data pipeline is deployed on a cloud server. This further facilitates a collaborative platform for different research groups regardless of their geographical locations. Based on the state-of-the-art in the Internet of Things (IoT) [47, 48], we have designed and implemented a robust and scalable real-time stream processing system leveraging Google Cloud infrastructure. The architecture is illustrated in **Sup. Fig. 9**. The IoT Core provides the functionalities to manage and configure connected devices conveniently and securely. Once the ECG signal is acquired from the measurement device, it is transmitted in real-time to the cloud platform through the Message Queuing Telemetry Transport (MQTT) protocol. The MQTT protocol is designed to maintain a long-lived connection between the device and the client with minimal communications overheads to save bandwidth for data transmission. Cloud Pub/Sub is an asynchronous communication medium between the device and the servers on the IoT cloud. The communication is based on the notion of topics that cache durable messages. Zebrafish ECG published on a certain topic by the device can be pushed to or pulled from the servers that subscribe to the same topic for storage and analytics.

The storage layer is responsible for storing the real-time zebrafish ECG into the database. As one of the most promising time-series databases, InfluxDB is employed in our proposed system for timely and reliable storage. As an advanced non-relational database, InfluxDB resolves the performance bottleneck of traditionally used databases such as MySQL, and provides greater flexibility and read-write speed. In the analytics layer, Cloud Functions, a serverless execution environment, run processing techniques such as denoising, filtering, normalizing, detecting P wave, QRS complex, or T wave, and extracting other useful features. Furthermore, machine learning approaches can be applied to these data with the aid of the Kubernetes Engine to train and deploy the models in containerized applications. Computationally intensive process and analysis tasks are carried out in powerful servers, which greatly eases the burden of local devices.

The graphical user interface is responsible for data visualization and management. It provides easy access to the data in the IoT cloud. Grafana is an open platform for monitoring time series data that we utilize to design dashboards to represent the ECG signals. Users can log onto the cloud to acquire visualized ECG data on either web pages or mobile applications. Based on the results of data analysis, we can observe and understand the real-time conditions of zebrafishes. In the event of any anomalies or suspicious readings, the IoT cloud will notify users in time.

The prolonged continuous ECG performance increases diagnostic and monitoring yield in the detection of asymptomatic cardiac events and reduces ECG artifact that improves arrhythmia detection. Moreover, it improves the quality and quantity of data collected from mutation-related sick sinus syndrome and arrhythmia during cardiac drug screening as well as increase zebrafish compliance provided by prolonged continuous ECG monitoring. However, we have also observed some limitations with our system. After conducting various experiments, we noticed that the electrode placement significantly contributes to the quality of ECG signal. With the use of the low Tricaine concentration to enable longer ECG recording, the fish tended to exhibit unexpected strong movement, thus leading to the electrode dislocation. Moreover, measuring multiple fish simultaneously required a great effort in aligning the electrode on the fish’s chest to acquire favorable ECG signal. The dripping setup used to continuously provide low Tricaine solution sometimes caused interferences to the ECG signal due to the inconsistency of its flow rate. To address these, we are planning to measure the real-time impedance of electrode-body interface and use that as the feedback information to control the electrode. We also plan to control the flow rate and provide a dedicated heating system to precisely control the medium temperature.

In conclusion, we have successfully demonstrated the novel Zebra II ECG system for multiple adult zebrafish. The major novelties lie in the long period (up to 1 hour) for recording of multiple fish, the minimal side effects, the automated cloud-based analytics as well as all other controlled features. We have demonstrated and further deployed the system for phenotyping cardiac mutants in response to various drugs and environmental cues treated simultaneously. Specifically, we have utilized our system to study Na^+^ sensitivity of the variant *SCN5A-D1275N*, for the first time, by observing changes in the frequency of sinus arrest episodes, heart rate value and QTc value. This is extremely important as it may provide answers for millions of cases of sudden death due to cardiac arrest. The Zebra II can be used for a host of cardiac disease studies and drug screening applications using the zebrafish model.

## Methods

### Mutant line *Tg(SCN5A-D1275N)* and zebrafish husbandry

Mutant line Tg(*SCN5A-D1275N*), a transgenic zebrafish arrhythmia model bearing the pathogenic human cardiac sodium channel mutation *SCN5A-D1275N*, was used to characterize and validate device performance, study sinus node dysfunction, and perform drug high throughput screening assays. Correlation between clinical phenotype and the mutant line has been reported for bradycardia, conduction-system abnormalities, episodes of sinus arrest [32].

Adult wild/mutant-type zebrafish with the age from 13 to 20 months (body lengths approximately 3-3.5 cm) were used in this study. Zebrafish are kept in a circulating system that continuously filters and aerates the system water to maintain the water quality required for a healthy aquatic environment. The fish room located in Engineering Gateway #3324 at UC Irvine is generally maintained between 26-28.5°C and the lighting conditions are controlled with 14:10 hours (*i.e.*, light: dark).

All animal protocols in this study were reviewed and approved by the Institutional Animal Care and Use Committee (IACUC) protocol (#AUP-18-115 at University of California, Irvine). All experiments were performed in accordance with relevant guidelines and regulations.

### Drug Administration

To anesthetize fish, we used a buffered solution of 200 parts-per-million (ppm) Tricaine (Sigma, USA) [49]. The Tricaine was dissolved in distilled water to a final concentration of 7,000 parts-per-million (ppm) as a stock, and the pH value was adjusted to 7.2 with sodium hydroxide (Sigma).

Amiodarone (Sigma) was dissolved in water at 65°C for 2 h and stocked as 900 μM at 4°C. Before use, the solution was re-dissolved at 65°C for 1 h [50]. The fish were immersed in a tank with 100 μM amiodarone for 1 hour and then return to fish water in 15 mins before doing experiment.

*SCN5A* is the gene associated with the alpha subunit of the voltage-gated sodium channel protein Nav1.5. Primarily responsible for the induction of the cardiac action potential, the Nav1.5 channel has been linked to several arrhythmogenic diseases such as sick sinus syndrome, long QT syndrome (LQTS), and Brugada syndrome [51]. Numerous studies have documented the molecular interactions and pathways involved with Nav1.5 as well as the various disease phenotypes exhibited by Nav1.5 variants [51, 52]. However, there is a current lack of functional characterization in regard to the molecular dynamics of Nav1.5 variants, including the functional response of Nav1.5 to initiate action potentials based on variation of sodium concentration. To the best of our knowledge, there are no research studies established to investigate the Na+ sensitivity in the development of sinus arrest. Therefore, an experiment was devised to determine the Na^+^ sensitivity of the variant *SCN5A-D1275N* by observing changes in the frequency of sinus arrest episodes, heart rate value and QTc value in the transgenic fish after treatment of 0.1, 0.3, 0.6, 0.9, and 1.8 ‰ of 5 M NaCl.

### Design of the Zebra II

The Zebra II comprises of a perfusion system, apparatuses, sensors and an in-house electronic system (**Fig. 1a**). The perfusion system comprises four syringes, four valves and tubing. Four syringes contain low dose Tricaine solution, continuously providing the solution to the fish through the tubing system to reduce the aggressiveness and activity of zebrafish but keep them awake. Four valves adjust the solution’s flow rate within a range of 5.5 - 6 ml/min [53]. Housing apparatuses and sensors are improved from the previous work [53]. Specifically, multiple small housings are made of polydimethylsiloxane (PDMS), providing comfort to the fish and thus minimizing unwanted artifacts. Moreover, the top and bottom of the apparatus are designed in such a way that the fish can lay comfortably on electrodes with curved shape in the bottom. The top has a redundant part sticking on the wall to keep the fish from escaping the apparatus. With the thermo box, a specific temperature is set by the thermostat control and the light bulb is turned on so that the box’s temperature can be maintained at the setup temperature and vice versa.

An in-house electronic system and a mobile application are described in **Fig. 1b-d, Sup. Fig. 1**. Specifically, the system includes a system on chip (SoC) supporting Bluetooth Low Energy (BLE), an analog front-end for zebrafish ECG signal acquisition, and a power-supply module with charge management. The SoC adopted nRF52832 from Nordic (*Trondheim, Norway*), which was a 64-MHz Arm Cortex-M4 CPU with a built-in BLE module. The analog front-end of ADS1299 from TI is well-known for biopotential measurements with eight low-noise, programmable gain amplifiers (PGAs) and eight high 24-bit resolution Delta-Sigma ADCs. Its high bit resolution provides both precision and dynamic range, allowing it to capture signals as high as 4.5 V and as low as 0.5 uV. The data rate is configurable from 250 samples per second (SPS) to 16 kSPS for all eight channels. Digital signals are sent from the ADS modules to the microcontroller for preprocessing via SPI interface. Data ready pin of ADS1299 module is triggered to signal microcontroller once a new data package is ready to transfer. Since multiple zebrafish are recorded simultaneously, differential mode configuration is utilized in the ADS1299. This would preclude signals in each fish from affecting each other. Moreover, a bias electrode with a signal generated by an ADS1299 chip is used, enabling a feedback loop built in the chip to get better common mode rejection.

### Investigation of tricaine and temperature to reduce cardiac rhythm side effect

As an important vertebrate model, zebrafish have been studied at both the embryonic and adult stages. Anesthesia is used in every experimental procedure to avoid discomfort, stress or pain. The most commonly used anesthetic drug is Tricaine (MS-222) and zebrafish usually are treated with 200 ppm in 3-5 minutes before experiment. However, this drug has been shown to affect cardiac function of the adult zebrafish and decrease the heart rate of the sedated subjects [21, 54]. As a result, MS-222 may skew the measurement of zebrafish physiological parameters. Moreover, temperature has profound effects on the performance and biogeography of ectothermic animals including fishes, partly through its effects on metabolism. At low temperatures (*e.g*., 18 - 20°C), myocyte activity is reduced as a natural adaptive mechanism to aid survival during colder climates or reasons [55], which leads to a reduction in heart rate. At higher temperatures, increased heart rate facilitates greater cardiac output to support a higher metabolic activity/demand for oxygen consistent with normal biological rate function. Therefore, with an optimal environment temperature, the proposed prolonged ECG system will help to lower the tricaine concentration which can reduce the cardiac rhythm side effect as well as maintain the ECG morphology. Three different Tricaine concentrations (*i.e.*, 75, 100, and 150 ppm) were used which account for 37.5%, 50% and 75% of original dose (200 ppm), respectively. Eight fish are used for each concentration group.

### Response analysis to drug treatment in real time with the Zebra II system

As a prominent vertebrae model for disease, zebrafish has already contributed to successful phenotype-based drug discovery, acting a bridge between in vitro assays and mammalian in vivo studies [56–58]. After establishing the optimized ECG assay conditions (*i.e.*, tricaine concentration, and temperature), the Zebra II demonstrated its potential in response to drug treatment in real time. The amiodarone drug was used in this experiment to demonstrate the drug screening capability of the system as it can affect ionic channels, associated with the potential for QT interval prolongation in the heart’s electric cycle. With different dosage (*i.e.*, 70 μM, 100 μM and 200 μM) consecutively treated for zebrafish, we can trigger the phenotype of QT interval prolongation to appear and vary in response to the drug treatment.

### Signal processing and statistics

The recorded ECG data were analyzed, and several parameters were extracted, including HR, QT, QTc intervals [49]. The standard deviation of normal sinus beats (SDNN) was used to evaluate the short term recording (5 minutes) beat-to-beat variance of HR and the standard deviation of the average normal-to-normal (NN) intervals for each of the 5 min segments (SDANN) was used for prolonged measurement (40 minutes) [59].

Statistical analysis was performed by using OriginLab 2019. Specifically, differences between samples were tested with student’s t-test and statistical significance accepted at a threshold of P < 0.05. Multiple comparisons were tested with one-way ANOVA and significant results (P < 0.05) were analyzed with pairwise comparisons using Student’s t-test applying significance levels adjusted with the Bonferroni method. Significant P-values are indicated with asterisks (*) with *P < 0.05, **P<0.01 and ***P <0.001. Correlation analysis was performed using Pearson’s correlation.

## Supporting information

shorturl.at/ctvzL

## Acknowledgement

The authors would like to acknowledge the financial support from the NSF CAREER Award #1917105 (H.C.), the NSF #1936519 (J.L. and H.C), the NIH SBIR grant #R44OD024874 (M.P.H.L and H.C.), the NIH HL107304 and HL081753 (X.X.). We thank Lauren Schmiess-Heine for managing the zebrafish facility; and staff members in the Xu lab at Mayo Clinic, MN, USA for maintaining and sharing the *Tg(SCN5A-D1275N)* fish.

## Author Contributions

### Conceptualization

Tai Le, Anh Hung Nguyen, Hung Cao.

### Data curation

Tai Le, Jimmy Zhang.

### Formal analysis

Tai Le, Jimmy Zhang, Anh Hung Nguyen.

### Investigation

Tai Le, Jimmy Zhang, Anh Hung Nguyen.

### Methodology

Tai Le, Jimmy Zhang, Anh Hung Nguyen, Ramses Trigo, Khuong Vo.

### Writing – original draft

Tai Le, Anh Hung Nguyen, Jimmy Zhang, Khuong Vo, Hung Cao.

### Writing – review & editing

Tai Le, Jimmy Zhang, Anh Hung Nguyen, Khuong Vo, Michael P.H. Lau, Juhyun Lee, Yonghe Ding, Xiaolei Xu, Hung Cao.

