## Supplementary material for "A Novel Art of Electrocardiogram Assessment in Zebrafish for Cardiovascular Disease Studies and Drug Screening": shorturl.at/ctvzL

### Supplementary document

(a) (b) (c)

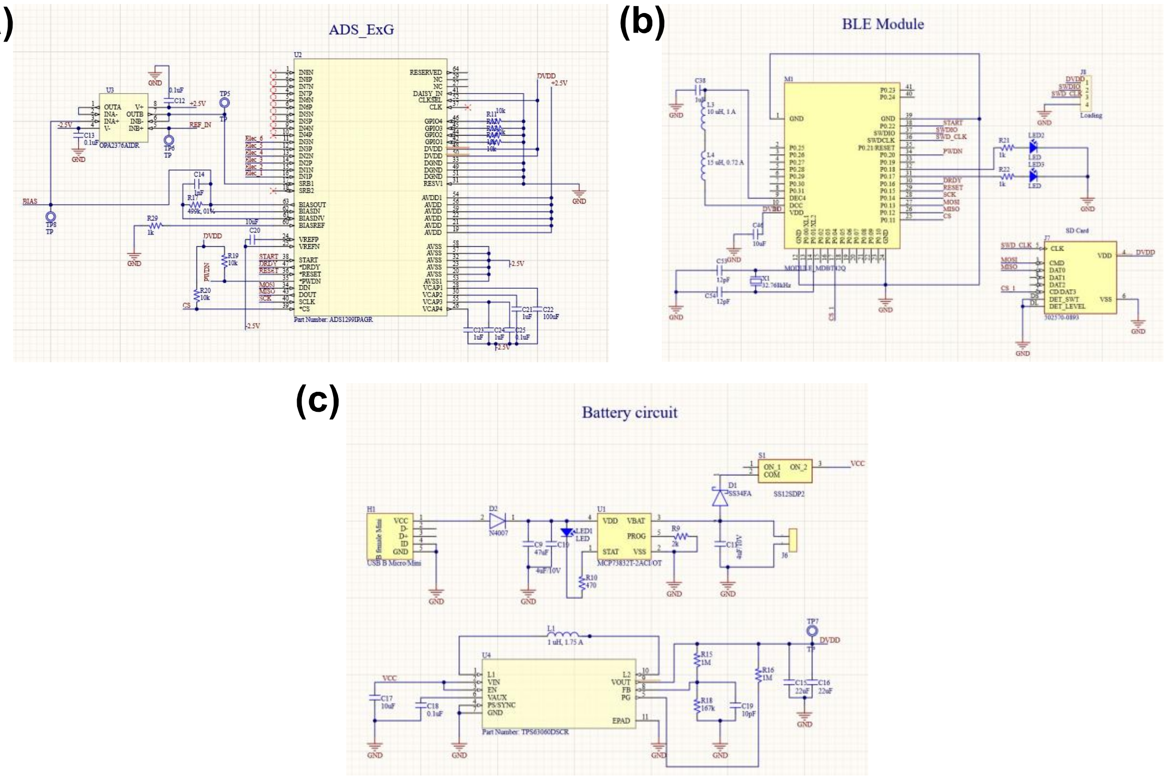


**Supplementary figure 1. Schematic of the ECG acquisition circuit.** (**a**) The analog front-end circuit, including ADS chip with 4 differential channels, an OP-AMP utilized to reduce common mode noise. (b) wireless transmission circuit, including a system-on-chip nRF52832. (c) The power-supply module included a charge management controller (MCP73832, *Microchip Inc*), a buck boost converter TPS63060 (*TI Instrumentation*) to maintain a 3.5 V output for the digital system regardless of the fluctuation of battery voltage level and two low-dropout regulators (i.e., TPS72325 and TPS73225) provide stable +- 2.5 V, respectively.

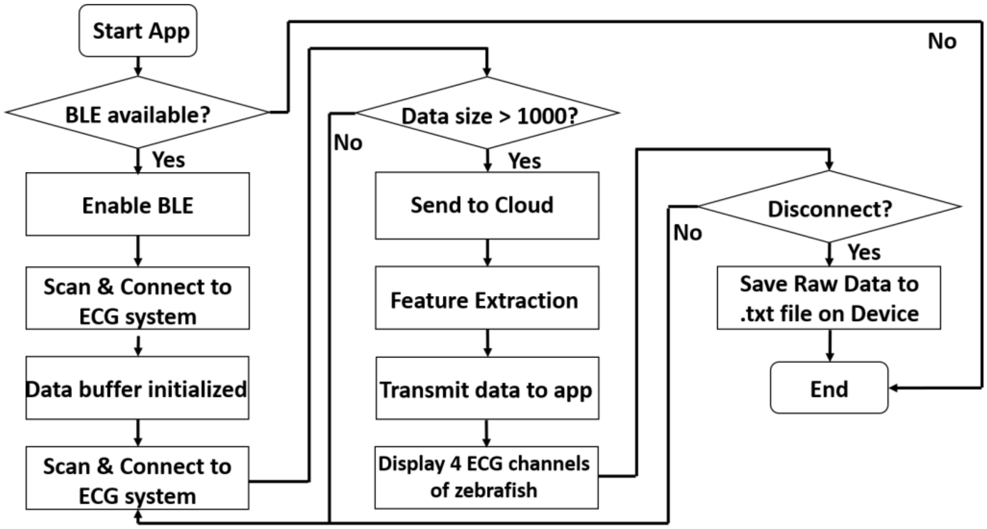


**Supplementary figure 2. Wireless data transmission to mobile application.** An Android application developed in Java that connects to the ECG system via BLE communication for data collection, displaying, and logging.

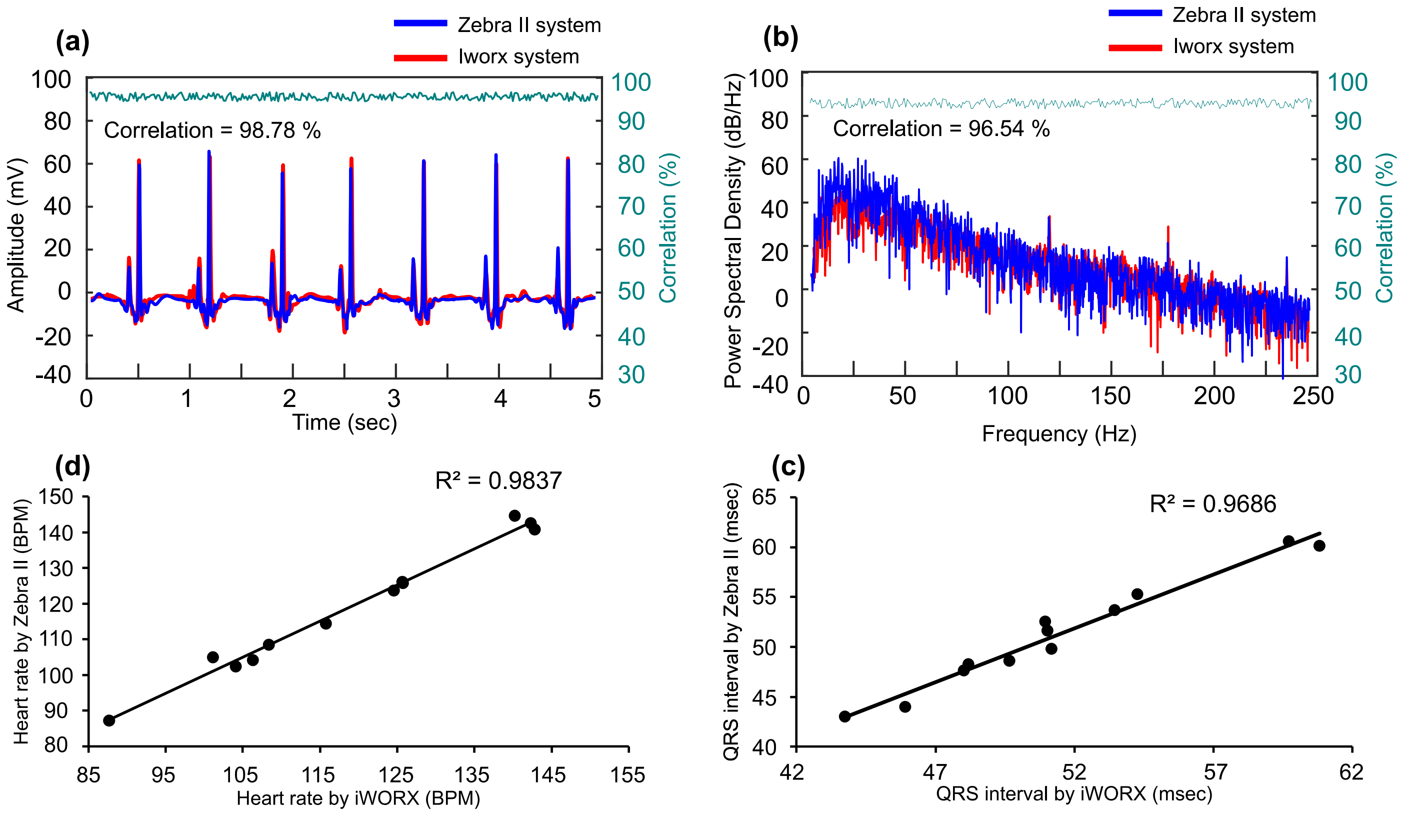


**Supplementary figure 3. Comparison of performance between Zebra II system and iWorx system.** The ECG measurement was conducted with two systems simultaneously (n = 8 fish). The collected data were then analyzed and compared in terms of correlation on time domain (**a**) and frequency domain (**b**). Correlation of heart rate (**d**) and QRS interval (**c**) extracted from ECG data collected by Zebra II and iWORX.

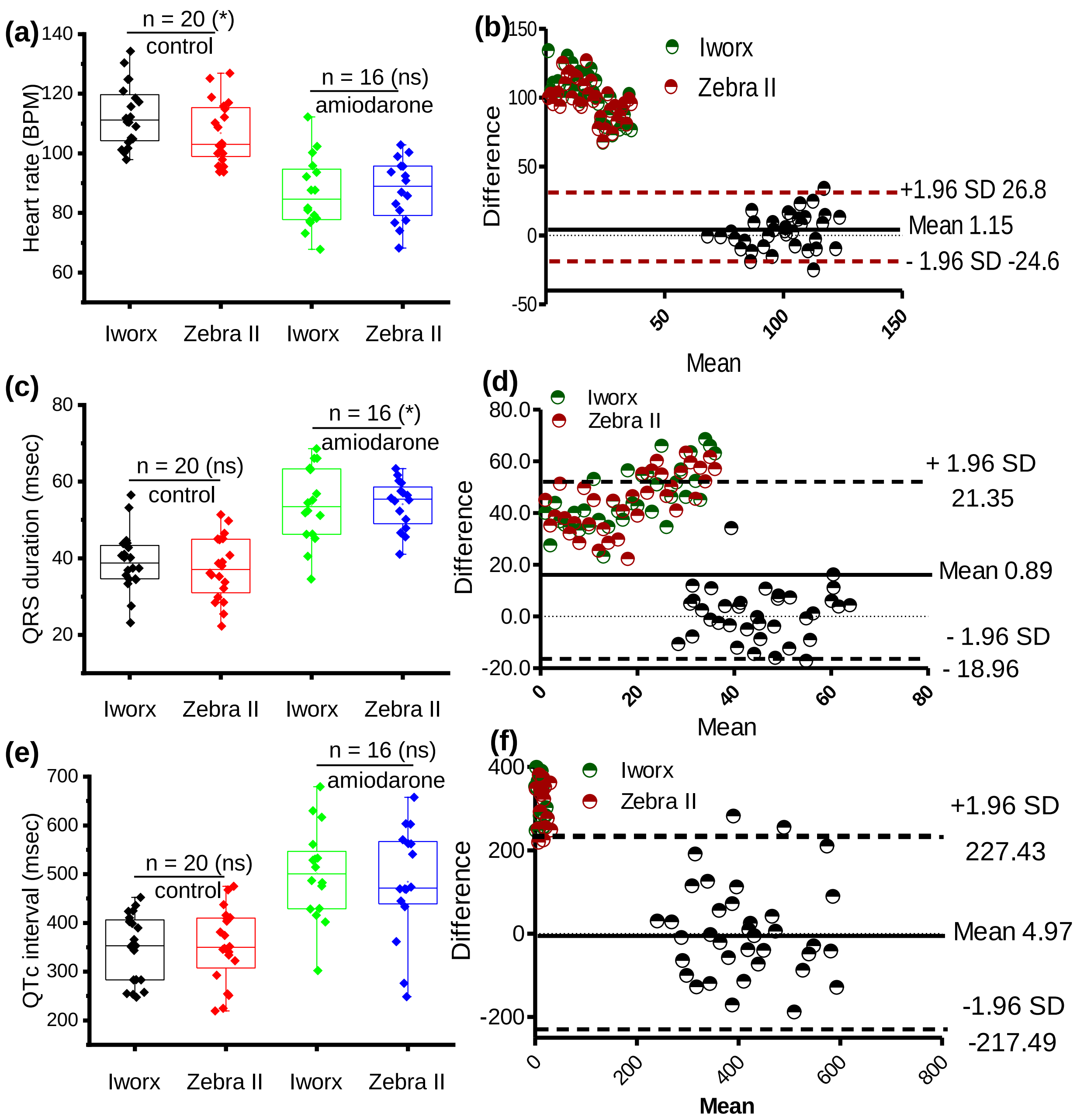


**Supplementary figure 4. Compare the performance between Zebra II and Iworx with amiodarone treatment.**

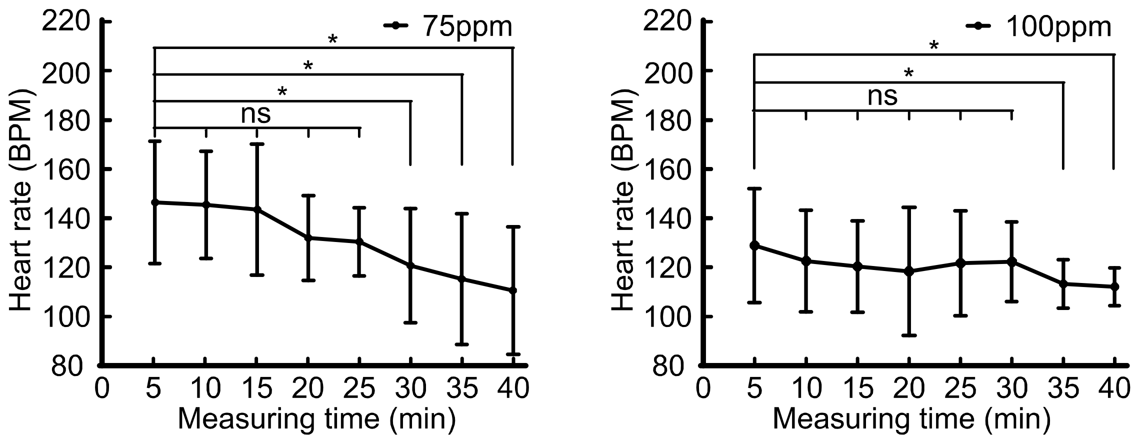


**Supplementary figure 5. Characterization of heart rate variation with different Tricaine concentration.** The experiment is conducted with n = 8 fish within 40-minute long ECG measurement, showing the heart rate variation with two Tricaine dosages in every 5 minutes.

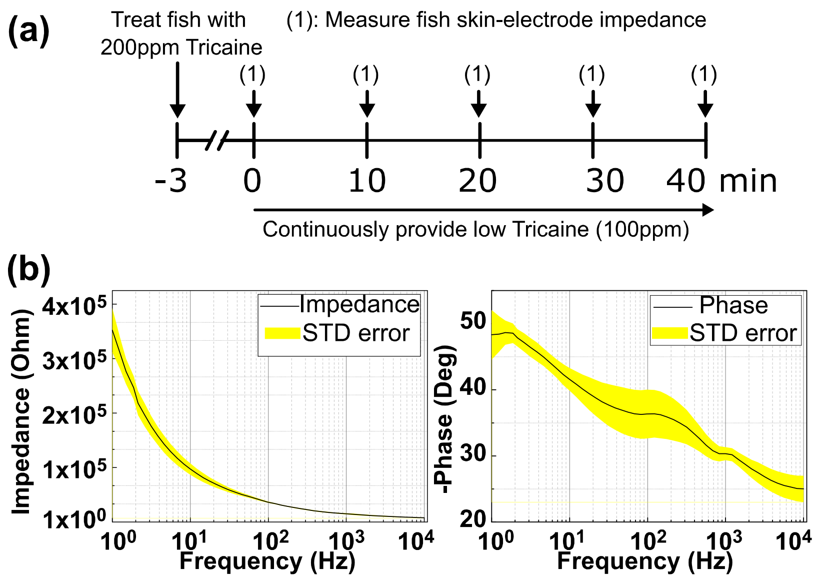


**Supplementary figure 6. Characterization of electrode-skin impedance.** (**a**) Timeline for experiment: Zebrafish were first anesthetized with 200ppm Tricaine in 3 minutes. Zebrafish were then loaded to each apparatus, followed by the tube system continuously providing low Tricaine concentration. The skin-electrode impedance was measured at 5 time points (i.e., every 10 minutes for 40 minutes). (**b**) Averaged impedance magnitude (left side) of 5 time points and phase (right side) of the electrodes placed on fish skin (n=8 animals).

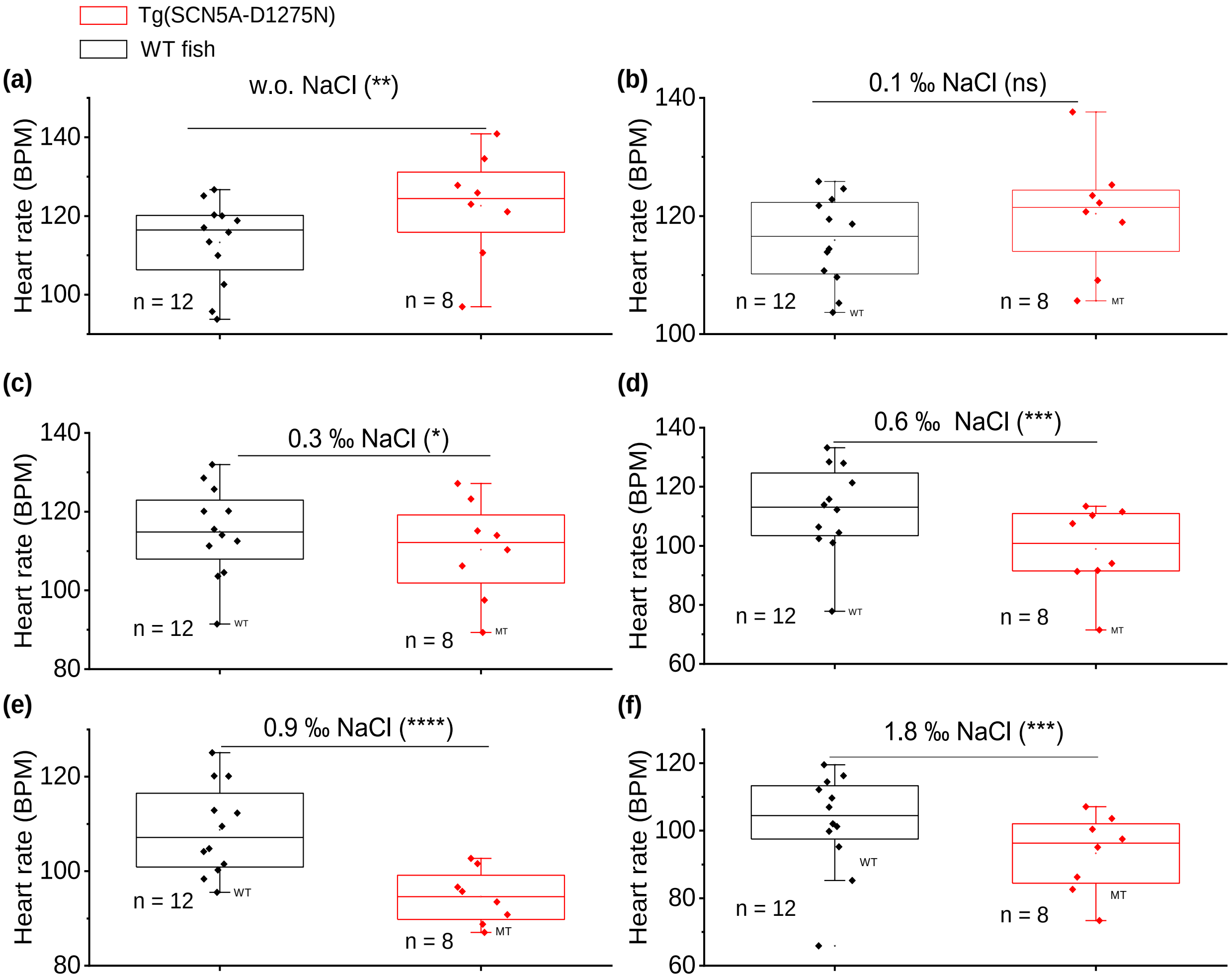


**Supplementary figure 7. Heart rate comparison between WT fish and Tg(SCN5A-D1275) with different NaCl concentrations.** Overall, the HR in WT fish does not show significant difference among NaCl concentrations. In contrast, the in mutant fish shows significant changes with the HR decreasing in response to the increase of NaCl dose.

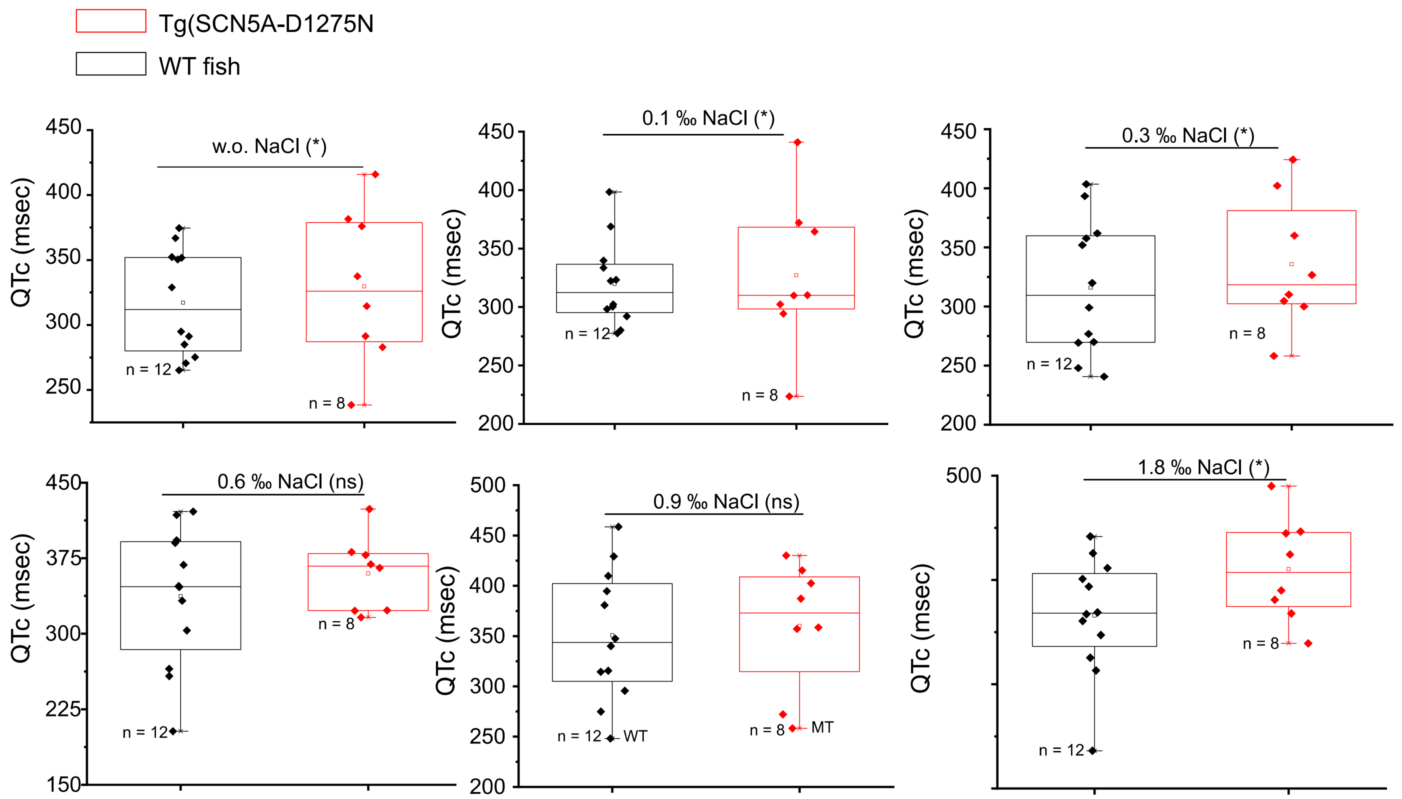


**Supplementary figure 8. QTc comparison between WT fish and Tg(SCN5A-D1275) with different NaCl concentration.** The significant difference showing in QTc value between two groups was found in 0.1, 0.3 and 1.8 ‰ NaCl. An increase in QTc value responding to the increase of NaCl level was found in both WT fish and mutant fish.

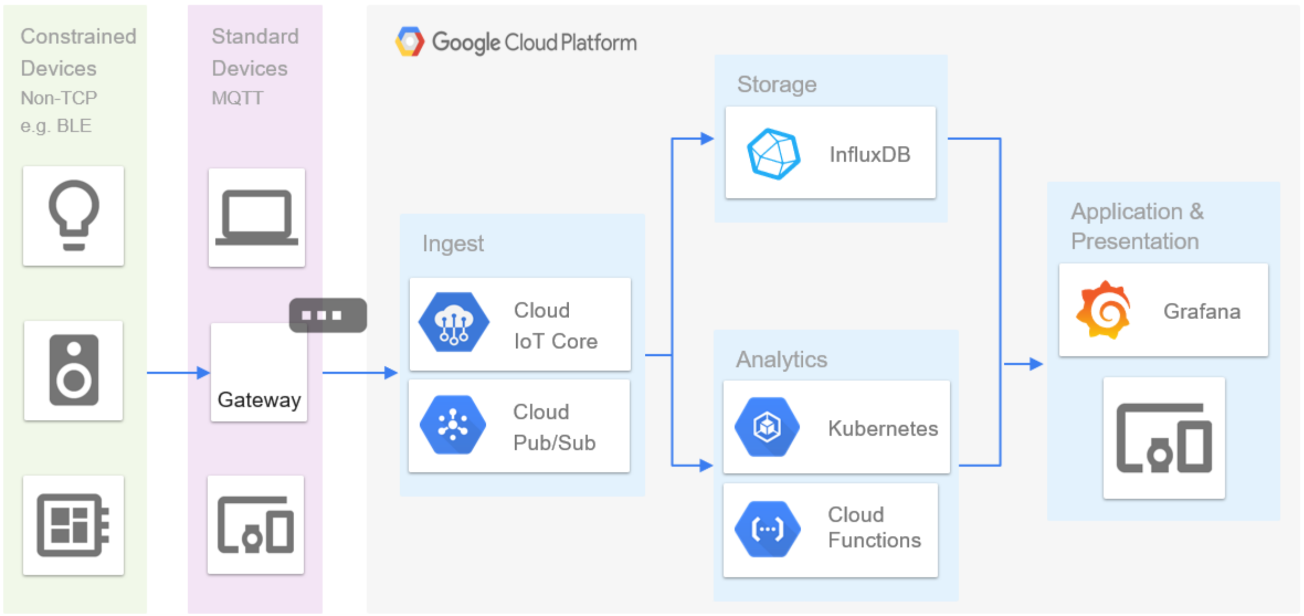


**Supplementary figure 9. The IOT system integrated with the prolonged ECG system.**

Table S1:

| Parameters | Numbers |
|---|---|
| Number of channels | 4 (can be up to 8) |
| Bandwidth | 1 Hz – 150 Hz |
| Input Range | ±2.5V |
| Resolution | 24 Bit |
| Sampling rate | 250 Hz |
| Common mode rejection ratio (CMRR © 60 Hz) | 100 dB |
| Input-Referred Noise | 1 µV |
| Enclosure | Plastic |
| Dimensions | 6.3’’W x 4.8’’L x 3.9’’H |
